# Characterization of velvet DNA-binding region by *Aspergillus nidulans* VelB

**DOI:** 10.64898/2026.06.04.730036

**Authors:** Wanping Chen, Anna M. Köhler, Gerhard H. Braus

**Affiliations:** Guangdong Provincial Key Laboratory of Microbial Culture Collection and Application, State Key Laboratory of Applied Microbiology Southern China, Institute of Microbiology, Guangdong Academy of Sciences, Guangzhou 510070, PR China; Department of Molecular Microbiology and Genetics, University of Göttingen, Göttingen 37077, Germany

**Author notes:** Corresponding Author: Wanping Chen.

**Keywords:** velvet regulators, VelB, DNA-binding, *Aspergillus*, fungi

## Abstract

Velvet regulators characterized by a conserved velvet domain function as a central hub that coordinately govern fungal development, secondary metabolism, stress adaptation, and pathogenicity. The velvet domain is organized into an N-terminal DNA-binding region of approximately 30 amino acids and a C-terminal dimerization region of roughly 100 amino acids. In this study, *Aspergillus nidulans* VelB was used as a paradigm to systematically dissect the velvet DNA-binding region. Three arginine residues R71, R80, and R81 in the N-terminal velvet domain indispensable for VelB function were identified through alanine-scanning mutagenesis of 15 conserved residues. Alanine substitutions at these positions caused severe defects in long-term spore viability, sexual development, and secondary metabolism. Further comparative characterization of electrostatic potential dynamics pre- and post-mutation revealed that the three individual substitutions markedly attenuate local electrostatic potential across the DNA-binding interface. Notably, these mutations drive comprehensive remodeling of the protein’s electrostatic properties, whereby electrostatic perturbations propagate across the entire protein exterior. Analysis of 4,999 velvet-domain sequences across the fungal kingdom revealed extraordinary conservation of these positions: arginine was present at position 71 in 85% of sequences, at position 80 in 91%, and at position 81 in 84%. Cross-kingdom complementation experiments further demonstrated that the wild-type velvet DNA-binding region from *Capsaspora owczarzaki*, a unicellular holozoan lacking the equivalent of R71, failed to rescue the *A. nidulans* velB deletion phenotype, whereas introduction of arginine at this position conferred substantial functional restoration. These findings establish that a cluster of conserved arginine residues generates the positive electrostatic surface potential required for velvet-DNA interaction, and define the molecular basis of DNA recognition by this ancient family of fungal transcription factors.

## Introduction

The coordination of morphological development with secondary metabolism represents a fundamental challenge in the life cycle of filamentous fungi (Calvo et al. 2002). These organisms must precisely orchestrate the timing of developmental transitions—such as the choice between asexual and sexual reproduction—with the production of bioactive secondary metabolites that serve critical functions in ecological competition, stress adaptation, and chemical defense (Bayram and Braus 2012; Liu et al. 2021). At the molecular level, this coordination is governed by a limited set of conserved transcriptional and epigenetic regulatory networks that integrate diverse environmental cues, including light, temperature, and nutrient availability, to direct appropriate cellular responses. Among these, the velvet family of transcriptional and epigenetic regulatory proteins stands out as a central hub that interconnects fungal development, secondary metabolism, stress adaptation, and pathogenicity, thereby functioning as a master orchestrator of fungal physiology across diverse fungal lineages (Chen et al. 2026).

The velvet regulatory system was first described over half a century ago in the model filamentous fungus *Aspergillus nidulans*, when Käfer (1965) identified a *veA* (velvet A) mutant strain (designated *veA1*) that exhibited a distinctive developmental phenotype (Kafer 1965). Unlike the wild-type strain, which favors asexual sporulation under illumination and sexual development in darkness, the *veA1* mutant displayed a marked shift toward constitutive asexual sporulation and severely impaired sexual development under normally permissive dark conditions, concomitantly losing the ability to produce major secondary metabolites (Kim et al. 2002; Stinnett et al. 2007). The *veA1* mutant colonies presented a characteristic velvet-like appearance due to their abundant asexual spore production, from which the "velvet" nomenclature was derived (Kafer 1965). Molecular characterization later revealed that the *veA1* allele carries a point mutation in the start codon that truncates the N-terminal 36 amino acids of VeA, thereby abolishing its nuclear localization signal and preventing light-regulated nuclear import (Stinnett et al. 2007).

Subsequent studies revealed that velvet proteins are highly conserved across the fungal kingdom, from early-diverging chytrids to advanced basidiomycetes, with homologs even identified beyond the fungal kingdom (Chen et al. 2024). The founding velvet family encompasses four core members: VeA, VelB (velvet-like B), VelC (velvet-like C), and VosA (viability of spores A) (Kim et al. 2002; Ni and Yu 2007; Park et al. 2012; Park et al. 2014). These proteins form multiple homomeric or heteromeric complexes in implementing a diverse range of functional roles, among which the trimeric velvet complex VelB–VeA–LaeA is a prominent example (Bayram et al. 2008; Strohdiek et al. 2025; Kohler et al. 2026). In darkness, VeA translocates into the nucleus and bridges VelB with the methyltransferase LaeA, thereby coordinating sexual development and secondary metabolite production. Under illumination, nuclear localization of VeA is reduced, resulting in altered complex assembly and a shift toward asexual development. This light-responsive regulatory mechanism represents one of the most important paradigms in fungal environmental adaptation.

At the structural level, all velvet proteins share a conserved velvet domain of approximately 200 amino acid residues (Chen et al. 2024). Crystallographic and molecular studies revealed that this domain exhibits structural similarity to the Rel homology domain of mammalian NF-κB transcription factors (Ahmed et al. 2013). The recent study proposed a general architecture of velvet domains consisting of an approximately 30-amino-acid N-terminal DNA-binding region and a roughly 100-amino-acid C-terminal dimerization region that contains α- and β-subunits separated by a flexible linker (Chen et al. 2025). These discoveries significantly advanced understanding of how velvet domains work.

At present, the C-terminal dimerization region has been functionally characterized (Chen et al. 2025), however, related studies on velvet DNA-binding region has not been reported. In this regard, the sequence features and key residues of velvet DNA-binding region were systemically investigated in this study by using *A. nidulans* VelB as a paradigm. Three key positive amino acid residues have been screened due to their indispensable roles for VelB function, which are prerequisites for creating positive electrostatic surface potential to facilitate interactions between velvet proteins and DNA.

## Materials and Methods

### *A. nidulans* and *Escherichia coli* strains and growth conditions

All *A. nidulans* strains applied and constructed in this work are summarized in Table 1. Strain AGB551 (*veA^+^*) served as the wild-type reference. Wild-type and mutant strains were cultured in minimal medium (MM). The medium consisted of 1% glucose, 7 mM KCl, 2 mM MgSO_4_, 70 mM NaNO_3_, 11.2 mM KH_2_PO_4_, and 0.1% trace element solution, with pH adjusted to 5.5, supplemented with 0.1% pyridoxine-HCl, 5 mM uridine, and 5 mM uracil (Barratt et al. 1965; Thieme et al. 2018). Strains were grown for 3 days on solid MM containing 2% agar at 37°C under constant illumination. Conidia re obtained by suspending spores from plate surfaces in sterile 0.96% NaCl solution containing 0.002% Tween 80, followed by removal of hyphal debris through Miracloth filtration. For phenotypic assessment of asexual development, approximately 200 spores were spotted onto the center of MM solid plates and incubated at 37°C under continuous light. For sexual development induction, plates were inoculated similarly, wrapped with Parafilm to prevent gas exchange, and incubated in complete darkness at 37°C.

**Table 1.**
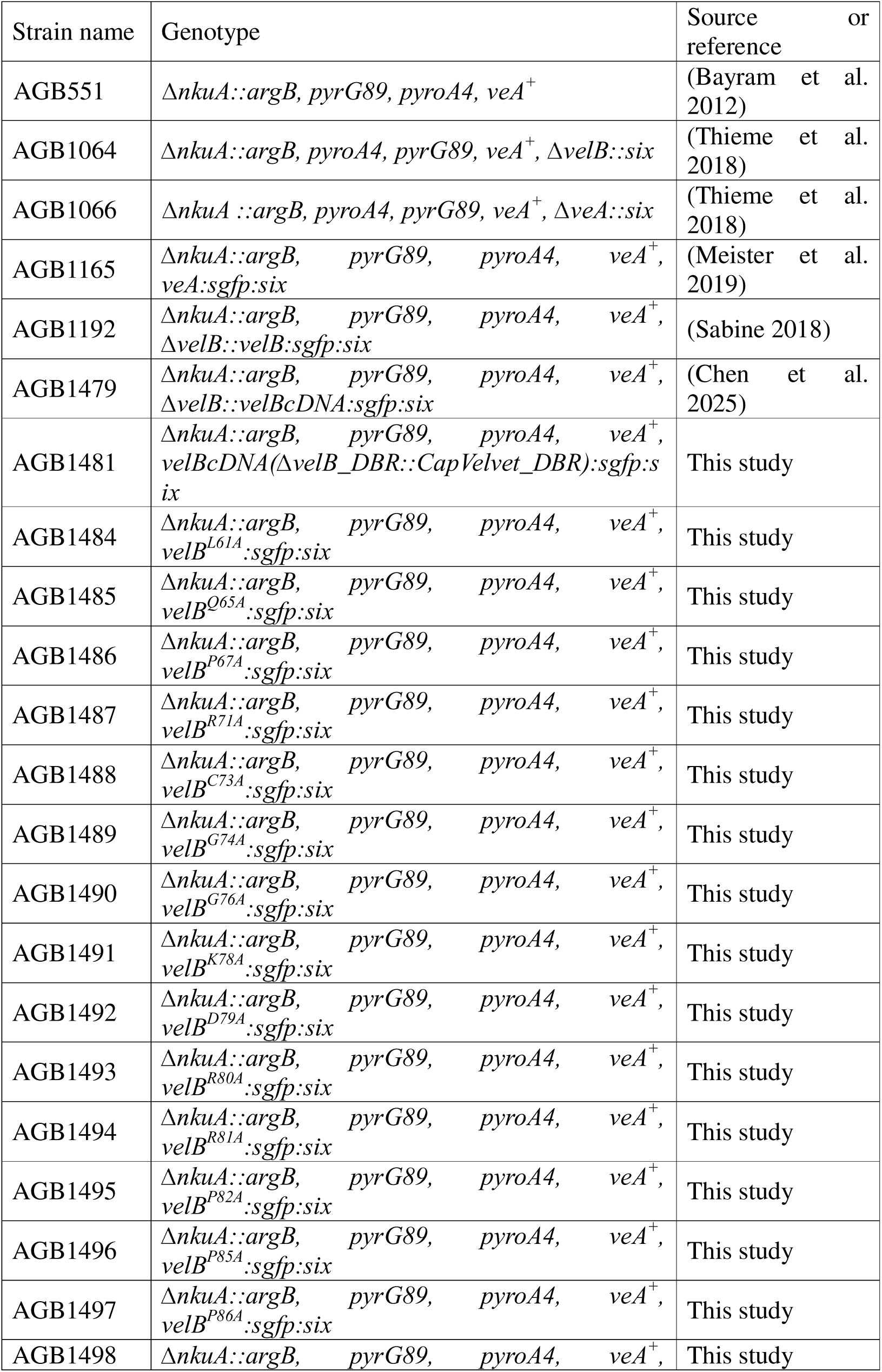
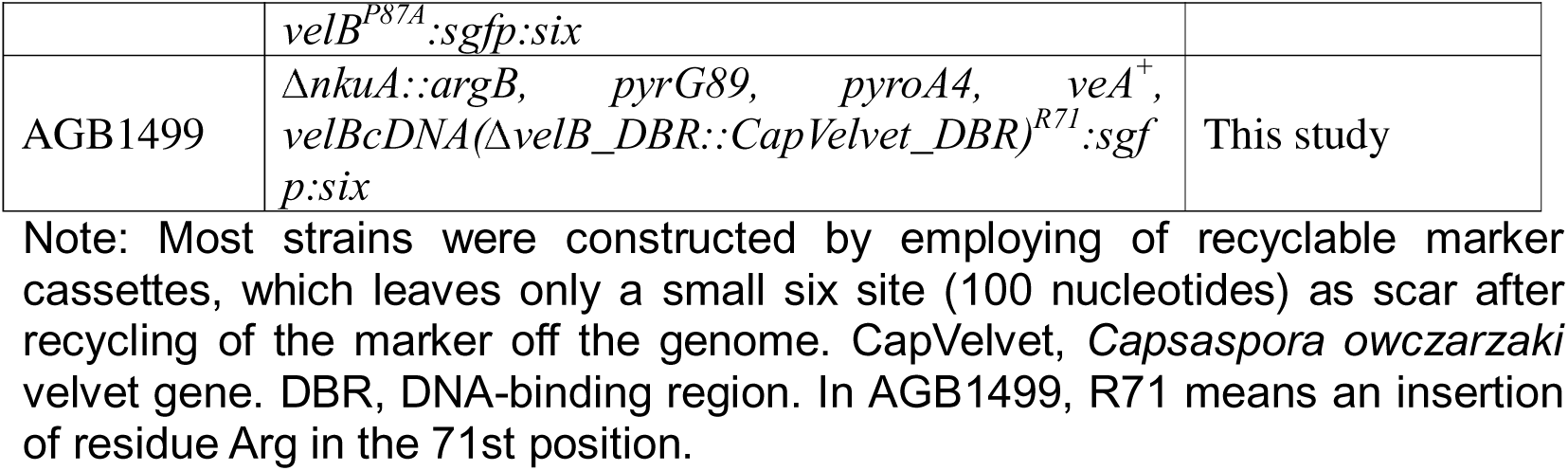
*A. nidulans* strains used in this study.

*E. coli* strains were grown on solid lysogeny broth (LB) medium (1% tryptone, 0.5% yeast extract, and 1% NaCl) or in liquid LB at 37°C with shaking at 200 rpm (Bertani 1951). Ampicillin (100 µg/mL) or kanamycin (50 µg/mL) was supplemented to avoid plasmid loss.

### Plasmid and strain preparation

Plasmids were amplified in *E. coli* and extracted using the Plasmid Spin Miniprep Kit (QIAGEN, Germany). Plasmid transformation into *E. coli* was carried out following established protocols (Inoue et al. 1990; Hanahan et al. 1991). Positive *E. coli* transformations on the selection medium were identified via colony polymerase chain reaction (PCR) and plasmid sequencing. All plasmids applied in the present study are summarized in Table S1. The oligonucleotides used in the construction of plasmids are listed in Table S2.

*A. nidulans* transformation was conducted via polyethylene glycol-mediated protoplast fusion according to published methods (Punt and van den Hondel 1992; Eckert et al. 2000). All transforming plasmids carrying recyclable phleomycin resistance cassettes are presented in Table S1. These plasmids were linearized by restriction digestion with PmeI, which possesses two recognition sites on each vector. The purified target fragments at a dosage of 5 μg were incubated with *A. nidulans* protoplasts. Successful genomic integration was validated by Southern blot hybridization using the AlkPhos Direct Labelling and Detection System (GE Healthcare Life Technologies, UK). All mutant strains generated herein are listed in **Table 1**.

### Construction of plasmid pME5335 and strains with replacement of *velB* DNA binding region by *C. owczarzaki* velvet one in *A*. *nidulans* (AGB1481)

The *velB* 5’ Untranslated Region (UTR) and 5’ fragment cloned with primers WC30 and WC154 from template pME5333, and *velB* cDNA remaining part fused with sgfp cloned from pME5333 with primers WC12 and WC155, were integrated into the SwaI restriction cutting site of plasmid pME5332 to generate pME5335, employing the Seamless Cloning and Assembly Kit (Invitrogen, Thermo Fisher Scientific, USA). The DNA sequence encoding the *C. owczarzaki* velvet DNA-binding region (amino acid sequence: LHIRQQPKHACMGGVSHDGARGNRR) was incorporated into the recombinant construct through primer-directed mutagenesis using primers WC154 and WC155. The linear *velBcDNA*(ΔvelB_DBR::CapVelvet_DBR):sgfp: phleoRM cassette excised from pME5335 by PmeI was integrated into AGB1064, resulting in AGB1481.

### Construction of plasmid pME5338 and *A*. *nidulans* mutant of *velB^L61A^* (AGB1484)

To generate the pME5338 recombinant plasmid, two targeted DNA fragments were amplified from the pME4687 template. The first fragment covering the *velB* 5′ UTR and its partial 5′ coding region was amplified using the primer pair WC30/WC208, while the second fragment containing the residual 3′ terminal sequence of *velB* fused to the *sgfp* reporter gene was amplified with WC12/WC209. These two fragments were seamlessly inserted into the SwaI restriction site of the pME5332 backbone utilizing a Seamless Cloning and Assembly Kit (Invitrogen, Thermo Fisher Scientific, USA). The L61A point mutation was incorporated into the *velB* coding region via primer-mediated mutagenesis using primers WC208 and WC209. The linear expression cassette *velB^L61A^: sgfp: phleoRM* was linearized from pME5338 through PmeI restriction digestion and transformed into the recipient *A. nidulans* strain AGB1064, yielding the final mutant strain AGB1484.

### Construction of plasmid pME5339 and *A*. *nidulans* mutant of *velB^Q65A^* (AGB1485)

To generate the pME5339 recombinant plasmid, two targeted DNA fragments were amplified from the pME4687 template. The first fragment covering the *velB* 5′ UTR and its partial 5′ coding region was amplified using the primer pair WC30/WC171, while the second fragment containing the residual 3′ terminal sequence of *velB* fused to the *sgfp* gene was amplified with WC12/WC172. These two fragments were seamlessly inserted into the SwaI restriction site of the pME5332 backbone utilizing a Seamless Cloning and Assembly Kit (Invitrogen, Thermo Fisher Scientific, USA). The Q65A point mutation was incorporated into the *velB* coding region via primer-mediated mutagenesis using primers WC171 and WC172. Subsequently, the linear expression cassette *velB^Q65A^: sgfp: phleoRM* was excised from pME5339 through PmeI restriction digestion and transformed into the recipient *A. nidulans* strain AGB1064, yielding the mutant strain AGB1485.

### Construction of plasmid pME5340 and *A*. *nidulans* mutant of *velB^P67A^* (AGB1486)

To generate the pME5340 recombinant plasmid, two targeted DNA fragments were amplified from the pME4687 template. The first fragment covering the *velB* 5′ UTR and its partial 5′ coding region was amplified using the primer pair WC30/WC173, while the second fragment containing the residual 3′ terminal sequence of *velB* fused to the *sgfp* gene was amplified with WC12/WC174. These two fragments were seamlessly inserted into the SwaI restriction site of the pME5332 backbone utilizing a Seamless Cloning and Assembly Kit (Invitrogen, Thermo Fisher Scientific, USA). The P67A point mutation was incorporated into the *velB* coding region via primer-mediated mutagenesis using primers WC173 and WC174. Subsequently, the linear expression cassette *velB^P67A^: sgfp: phleoRM* was excised from pME5340 through PmeI restriction digestion and transformed into the recipient *A. nidulans* strain AGB1064, yielding the mutant strain AGB1486.

### Construction of plasmid pME5341 and *A*. *nidulans* mutant of *velB^R71A^* (AGB1487)

To generate the pME5341 recombinant plasmid, two targeted DNA fragments were amplified from the pME4687 template. The first fragment covering the *velB* 5′ UTR and its partial 5′ coding region was amplified using the primer pair WC30/WC175, while the second fragment containing the residual 3′ terminal sequence of *velB* fused to the *sgfp* gene was amplified with WC12/WC176. These two fragments were seamlessly inserted into the SwaI restriction site of the pME5332 backbone utilizing a Seamless Cloning and Assembly Kit (Invitrogen, Thermo Fisher Scientific, USA). The R71A point mutation was incorporated into the *velB* coding region via primer-mediated mutagenesis using primers WC175 and WC176. Subsequently, the linear expression cassette *velB^R71A^: sgfp: phleoRM* was excised from pME5341 through PmeI restriction digestion and transformed into the recipient *A. nidulans* strain AGB1064, yielding the mutant strain AGB1487.

### Construction of plasmid pME5342 and *A*. *nidulans* mutant of *velB^C73A^* (AGB1488)

To generate the pME5342 recombinant plasmid, two targeted DNA fragments were amplified from the pME4687 template. The first fragment covering the *velB* 5′ UTR and its partial 5′ coding region was amplified using the primer pair WC30/WC251, while the second fragment containing the residual 3′ terminal sequence of *velB* fused to the *sgfp* gene was amplified with WC12/WC252. These two fragments were seamlessly inserted into the SwaI restriction site of the pME5332 backbone utilizing a Seamless Cloning and Assembly Kit (Invitrogen, Thermo Fisher Scientific, USA). The C73A point mutation was incorporated into the *velB* coding region via primer-mediated mutagenesis using primers WC251 and WC252. Subsequently, the linear expression cassette *velB^C73A^: sgfp: phleoRM* was excised from pME5342 through PmeI restriction digestion and transformed into the recipient *A. nidulans* strain AGB1064, yielding the mutant strain AGB1488.

### Construction of plasmid pME5343 and *A*. *nidulans* mutant of *velB^G74A^* (AGB1489)

To generate the pME5343 recombinant plasmid, two targeted DNA fragments were amplified from the pME4687 template. The first fragment covering the *velB* 5′ UTR and its partial 5′ coding region was amplified using the primer pair WC30/WC177, while the second fragment containing the residual 3′ terminal sequence of *velB* fused to the *sgfp* gene was amplified with WC12/WC178. These two fragments were seamlessly inserted into the SwaI restriction site of the pME5332 backbone utilizing a Seamless Cloning and Assembly Kit (Invitrogen, Thermo Fisher Scientific, USA). The G74A point mutation was incorporated into the *velB* coding region via primer-mediated mutagenesis using primers WC177 and WC178. Subsequently, the linear expression cassette *velB^G74A^: sgfp: phleoRM* was excised from pME5343 through PmeI restriction digestion and transformed into the recipient *A. nidulans* strain AGB1064, yielding the mutant strain AGB1489.

### Construction of plasmid pME5344 and *A*. *nidulans* mutant of *velB^G76A^* (AGB1490)

To generate the pME5344 recombinant plasmid, two targeted DNA fragments were amplified from the pME4687 template. The first fragment covering the *velB* 5′ UTR and its partial 5′ coding region was amplified using the primer pair WC30/WC210, while the second fragment containing the residual 3′ terminal sequence of *velB* fused to the *sgfp* gene was amplified with WC12/WC211. These two fragments were seamlessly inserted into the SwaI restriction site of the pME5332 backbone utilizing a Seamless Cloning and Assembly Kit (Invitrogen, Thermo Fisher Scientific, USA). The G76A point mutation was incorporated into the *velB* coding region via primer-mediated mutagenesis using primers WC210 and WC211. Subsequently, the linear expression cassette *velB^G76A^: sgfp: phleoRM* was excised from pME5344 through PmeI restriction digestion and transformed into the recipient *A. nidulans* strain AGB1064, yielding the mutant strain AGB1490.

### Construction of plasmid pME5345 and *A*. *nidulans* mutant of *velB^K78A^* (AGB1491)

To generate the pME5345 recombinant plasmid, two targeted DNA fragments were amplified from the pME4687 template. The first fragment covering the *velB* 5′ UTR and its partial 5′ coding region was amplified using the primer pair WC30/WC179, while the second fragment containing the residual 3′ terminal sequence of *velB* fused to the *sgfp* gene was amplified with WC12/WC180. These two fragments were seamlessly inserted into the SwaI restriction site of the pME5332 backbone utilizing a Seamless Cloning and Assembly Kit (Invitrogen, Thermo Fisher Scientific, USA). The K78A point mutation was incorporated into the *velB* coding region via primer-mediated mutagenesis using primers WC179 and WC180. Subsequently, the linear expression cassette *velB^K78A^: sgfp: phleoRM* was excised from pME5345 through PmeI restriction digestion and transformed into the recipient *A. nidulans* strain AGB1064, yielding the mutant strain AGB1491.

### Construction of plasmid pME5346 and *A*. *nidulans* mutant of *velB^D79A^* (AGB1492)

To generate the pME5346 recombinant plasmid, two targeted DNA fragments were amplified from the pME4687 template. The first fragment covering the *velB* 5′ UTR and its partial 5′ coding region was amplified using the primer pair WC30/WC181, while the second fragment containing the residual 3′ terminal sequence of *velB* fused to the *sgfp* gene was amplified with WC12/WC182. These two fragments were seamlessly inserted into the SwaI restriction site of the pME5332 backbone utilizing a Seamless Cloning and Assembly Kit (Invitrogen, Thermo Fisher Scientific, USA). The D79A point mutation was incorporated into the *velB* coding region via primer-mediated mutagenesis using primers WC181 and WC182. Subsequently, the linear expression cassette *velB^D79A^: sgfp: phleoRM* was excised from pME5346 through PmeI restriction digestion and transformed into the recipient *A. nidulans* strain AGB1064, yielding the mutant strain AGB1492.

### Construction of plasmid pME5347 and *A*. *nidulans* mutant of *velB^R80A^* (AGB1493)

To generate the pME5347 recombinant plasmid, two targeted DNA fragments were amplified from the pME4687 template. The first fragment covering the *velB* 5′ UTR and its partial 5′ coding region was amplified using the primer pair WC30/WC183, while the second fragment containing the residual 3′ terminal sequence of *velB* fused to the *sgfp* gene was amplified with WC12/WC184. These two fragments were seamlessly inserted into the SwaI restriction site of the pME5332 backbone utilizing a Seamless Cloning and Assembly Kit (Invitrogen, Thermo Fisher Scientific, USA). The R80A point mutation was incorporated into the *velB* coding region via primer-mediated mutagenesis using primers WC183 and WC184. Subsequently, the linear expression cassette *velB^R80A^: sgfp: phleoRM* was excised from pME5347 through PmeI restriction digestion and transformed into the recipient *A. nidulans* strain AGB1064, yielding the mutant strain AGB1493.

### Construction of plasmid pME5348 and *A*. *nidulans* mutant of *velB^R81A^* (AGB1494)

To generate the pME5348 recombinant plasmid, two targeted DNA fragments were amplified from the pME4687 template. The first fragment covering the *velB* 5′ UTR and its partial 5′ coding region was amplified using the primer pair WC30/WC185, while the second fragment containing the residual 3′ terminal sequence of *velB* fused to the *sgfp* gene was amplified with WC12/WC186. These two fragments were seamlessly inserted into the SwaI restriction site of the pME5332 backbone utilizing a Seamless Cloning and Assembly Kit (Invitrogen, Thermo Fisher Scientific, USA). The R81A point mutation was incorporated into the *velB* coding region via primer-mediated mutagenesis using primers WC185 and WC186. Subsequently, the linear expression cassette *velB^R81A^: sgfp: phleoRM* was excised from pME5348 through PmeI restriction digestion and transformed into the recipient *A. nidulans* strain AGB1064, yielding the mutant strain AGB1494.

### Construction of plasmid pME5349 and *A*. *nidulans* mutant of *velB^P82A^* (AGB1495)

To generate the pME5349 recombinant plasmid, two targeted DNA fragments were amplified from the pME4687 template. The first fragment covering the *velB* 5′ UTR and its partial 5′ coding region was amplified using the primer pair WC30/WC187, while the second fragment containing the residual 3′ terminal sequence of *velB* fused to the *sgfp* gene was amplified with WC12/WC188. These two fragments were seamlessly inserted into the SwaI restriction site of the pME5332 backbone utilizing a Seamless Cloning and Assembly Kit (Invitrogen, Thermo Fisher Scientific, USA). The P82A point mutation was incorporated into the *velB* coding region via primer-mediated mutagenesis using primers WC187 and WC188. Subsequently, the linear expression cassette *velB^P82A^: sgfp: phleoRM* was excised from pME5349 through PmeI restriction digestion and transformed into the recipient *A. nidulans* strain AGB1064, yielding the mutant strain AGB1495.

### Construction of plasmid pME5350 and *A*. *nidulans* mutant of *velB^P85A^* (AGB1496)

To generate the pME5350 recombinant plasmid, two targeted DNA fragments were amplified from the pME4687 template. The first fragment covering the *velB* 5′ UTR and its partial 5′ coding region was amplified using the primer pair WC30/WC189, while the second fragment containing the residual 3′ terminal sequence of *velB* fused to the *sgfp* gene was amplified with WC12/WC190. These two fragments were seamlessly inserted into the SwaI restriction site of the pME5332 backbone utilizing a Seamless Cloning and Assembly Kit (Invitrogen, Thermo Fisher Scientific, USA). The P85A point mutation was incorporated into the *velB* coding region via primer-mediated mutagenesis using primers WC189 and WC190. Subsequently, the linear expression cassette *velB^P85A^: sgfp: phleoRM* was excised from pME5350 through PmeI restriction digestion and transformed into the recipient *A. nidulans* strain AGB1064, yielding the mutant strain AGB1496.

### Construction of plasmid pME5351 and *A*. *nidulans* mutant of *velB^P86A^* (AGB1497)

To generate the pME5351 recombinant plasmid, two targeted DNA fragments were amplified from the pME4687 template. The first fragment covering the *velB* 5′ UTR and its partial 5′ coding region was amplified using the primer pair WC30/WC191, while the second fragment containing the residual 3′ terminal sequence of *velB* fused to the *sgfp* gene was amplified with WC12/WC192. These two fragments were seamlessly inserted into the SwaI restriction site of the pME5332 backbone utilizing a Seamless Cloning and Assembly Kit (Invitrogen, Thermo Fisher Scientific, USA). The P86A point mutation was incorporated into the *velB* coding region via primer-mediated mutagenesis using primers WC191 and WC192. Subsequently, the linear expression cassette *velB^P86A^: sgfp: phleoRM* was excised from pME5351 through PmeI restriction digestion and transformed into the recipient *A. nidulans* strain AGB1064, yielding the mutant strain AGB1497.

### Construction of plasmid pME5352 and *A*. *nidulans* mutant of *velB^P87A^* (AGB1498)

To generate the pME5352 recombinant plasmid, two targeted DNA fragments were amplified from the pME4687 template. The first fragment covering the *velB* 5′ UTR and its partial 5′ coding region was amplified using the primer pair WC30/WC193, while the second fragment containing the residual 3′ terminal sequence of *velB* fused to the *sgfp* gene was amplified with WC12/WC194. These two fragments were seamlessly inserted into the SwaI restriction site of the pME5332 backbone utilizing a Seamless Cloning and Assembly Kit (Invitrogen, Thermo Fisher Scientific, USA). The P87A point mutation was incorporated into the *velB* coding region via primer-mediated mutagenesis using primers WC193 and WC194. Subsequently, the linear expression cassette *velB^P87A^: sgfp: phleoRM* was excised from pME5352 through PmeI restriction digestion and transformed into the recipient *A. nidulans* strain AGB1064, yielding the mutant strain AGB1498.

### Construction of plasmid pME5353 and *A*. *nidulans* mutant of *velBcDNA(*Δ*velB_DBR::CapVelvet_DBR)^R71^* (AGB1499)

The *velB* 5’ UTR and 5’ fragment cloned with primers WC30 and WC212, and the remaining 3’ fragment fused with sgfp cloned with primers WC12 and WC213 from template pME5335 were integrated into the SwaI restriction cutting site of plasmid pME5332 to generate pME5353, employing the Seamless Cloning and Assembly Kit (Invitrogen, Thermo Fisher Scientific, USA). The mutant *velBcDNA(*Δ*velB_DBR::CapVelvet_DBR)^R71^*was introduced by primers WC212 and WC213. The linear *velBcDNA(*Δ*velB_DBR::CapVelvet_DBR)^R71^*: *sgfp*: phleoRM cassette from the digestion of pME5353 by PmeI was integrated into AGB1064, resulting in AGB1499.

### Sequence logo creation of DNA-binding domains

Multiple protein sequence alignments corresponding to the DNA-binding domains of six transcription factor families, including Rel (PF00554), zinc finger (PF01530), homeobox (PF00046), basic region leucine zipper (PF00170), SRF-TF (PF00319), and helix-loop-helix (PF00010), were retrieved from the InterPro database (Paysan-Lafosse et al. 2024; Blum et al. 2025). For the velvet DNA-binding domain, sequence alignment of 4,999 putative velvet domains was constructed in accordance with a previously reported protocol (Chen et al. 2025). Conserved sequence logos based on the aforementioned alignments were generated using WebLogo 3 (Crooks et al. 2004). Subsequent sequence editing and statistics analyses were implemented via Jalview Desktop (version 2.11.5.1) (Waterhouse et al. 2009).

### Protein isolation of *A. nidulans* mycelia

Protein extraction from *A. nidulans* mycelia was conducted following a previously described protocol (Thieme et al. 2018). Briefly, *A. nidulans* strains were cultivated vegetatively in 500mL MM inoculated with 5x10^8^ spores at 37°C for 20 h. The harvested mycelia were collected using sterile filters (Merck, Darmstadt, Germany), rinsed with saline-PMSF-DMSO, dehydrated, and immediately snap-frozen in liquid nitrogen. Frozen mycelial samples were homogenized with a table mill under liquid nitrogen conditions. The pulverized tissues were resuspended in an equal volume of B+ buffer (300 mM NaCl, 100 mM Tris pH 7.5, 10% glycerol, 1 mM EDTA, 0.1% NP-40). The buffer was pre-supplemented with 1.5 mM DTT, complete EDTA-free protease inhibitor cocktail (ROCHE Diagnostics GmbH, Basel, Switzerland), and 0.001 mM PMSF. The resulting homogenate was centrifuged at 13,000 rpm for 30 min at 4 °C. The clarified supernatant was aliquoted into new centrifuge tubes and stored at −20 °C. The protein concentration of each sample was quantified using a NanoDrop ND-1000 spectrophotometer.

### Western blot analysis of *A. nidulans* protein extracts

GFP fusion proteins was detected via Western blotting based on a previously established method (Chen et al. 2025). Briefly, crude protein samples were denatured with 3x protein loading buffer composed of 250 mM Tris-HCl (pH 6.8), 15% (v/v) β-mercaptoethanol, 30% (v/v) glycerol, 7% (v/v) SDS, and 0.3% (w/v) bromophenol blue. Protein samples were subjected to thermal denaturation at 95 °C for 5 minutes. Equal amounts of denatured crude protein extracts (90 µg per lane) from each fungal strain were resolved by sodium dodecyl sulfate-polyacrylamide gel electrophoresis (SDS-PAGE) using 12% polyacrylamide gels. Separated proteins were electrophoretically transferred onto nitrocellulose membranes (Merck, Darmstadt, Germany) at a constant voltage of 100 V for 1 hour.

Membranes were blocked with 5% (w/v) skim milk powder dissolved in Tris-buffered saline containing 0.05% Tween 20 (TBST; 10 mM Tris-HCl, pH 8.0, 150 mM NaCl) for 1 hour at room temperature. The blocked membranes were then incubated with a mouse monoclonal anti-GFP primary antibody (sc-9996, Santa Cruz Biotechnology) diluted 1:250 in blocking buffer. Following three consecutive washes with TBST, membranes were incubated with a horseradish peroxidase-conjugated goat anti-mouse secondary antibody (115-035-003, Jackson Immuno Research, West Grove, USA) at a 1:1000 dilution for 1 hour with gentle agitation at room temperature. Membranes were subsequently washed three additional times with TBST to remove unbound secondary antibody.

Chemiluminescent detection was performed using an enhanced chemiluminescence (ECL) reagent prepared immediately prior to use. Solution A consisted of 9 mL deionized water, 1 mL 1 M Tris-HCl (pH 8.5), 100 µL 250 mM luminol, and 44 µL 400 mM p-coumaric acid. Solution B contained 9 mL deionized water, 1 mL 1 M Tris-HCl (pH 8.5), and 6.14 µL 30% hydrogen peroxide. Equal volumes of Solution A and Solution B were mixed and applied to the membrane, which was then incubated for 2 minutes in the dark with gentle rocking. Chemiluminescent signals were captured using the Fusion-SL7 imaging system (Vilber Lourmat) and quantified using Fusion software (version 15.18) and Bio1D software (version 15.08).

### Extraction of secondary metabolites and their LC-MS analysis

Extraction of secondary metabolites and subsequent LC-MS analysis were performed as described previously (Chen et al. 2025). Briefly, each fungal strain was inoculated with 1×10D fresh spores on solid MM supplemented with 0.1% pyridoxine-HCl, 5 mM uridine, and 5 mM uracil, and incubated in the dark at 37 °C for 7 days. Metabolites were extracted from two colony agar plugs by homogenization in 5 mL water and 5 mL ethyl acetate, followed by overnight shaking at 220 rpm. After centrifugation at 2500 rpm for 5 min, the ethyl acetate phase was collected, evaporated to dryness, and stored at -20 °C. All extractions were performed in triplicate.

For LC-MS analysis, extracts were reconstituted in 500 µL acetonitrile/water (1:1) and centrifuged at 13,000 rpm for 10 min at 4 °C. Chromatographic separation was performed on an Acclaim 120 C18 column (4.6 × 100 mm, 5 µm) using a 0.1% formic acid/acetonitrile gradient (5–95% B in 20 min, 10 min isocratic wash) at 0.8 mL/min and 30 °C. Mass spectra were acquired in positive and negative ESI modes (m/z 70–1050) on a Q Exactive Focus Orbitrap mass spectrometer coupled to an UltiMate 3000 HPLC system. Data were analyzed using Xcalibur 4.4 and FreeStyle 1.8 SP2 software.

### Conidiospore and cleistothecia quantification

Fresh conidia were harvested from 3-day-old fungal cultures grown on solid MM plates using sterile 0.96% (w/v) NaCl solution supplemented with 0.002% (v/v) Tween 80. The resulting conidial suspensions were filtered through sterile Miracloth to remove residual mycelial debris, and the filtrates were used as conidial stock solutions. Stock suspensions were diluted to the required concentrations with sterile distilled water as needed. Conidial concentrations were quantified using a Coulter Z2 particle counter (Beckman Coulter GmbH, Krefeld, Germany).

Cleistothecia quantification was performed as described previously (Thieme et al. 2018). Briefly, 5 mm-diameter agar plugs (excised using the wide end of a standard 200 μL pipette tip) were harvested from representative regions of mature colonies. Cleistothecia were gently teased apart and dispersed onto a fresh sterile agar plate to ensure complete separation of individual fruiting bodies. Cleistothecial counts were determined using an SZX12-ILLB2-200 stereomicroscope (Olympus Corporation, Tokyo, Japan). At least five independent agar plugs were analyzed per strain, and all experiments were performed in biological triplicate.

### Spore viability assay

Spore viability assays were performed as described previously (Chen et al. 2025). Briefly, fresh conidia were harvested from 3-day-old fungal cultures grown on solid minimal medium (MM) agar plates, and resuspended in sterile 0.96% (w/v) NaCl solution supplemented with 0.002% (v/v) Tween 80. Conidial suspensions were filtered through sterile Miracloth to remove mycelial debris, and serially diluted to a final concentration of 1×10D conidia/mL. Stock suspensions were stored at 4 °C in the dark until use.

For viability assessment at designated time points post-harvest, stock spore solutions were further serially diluted. Aliquots (100 μL) containing approximately 100 conidia were spread evenly onto solid MM agar plates, and incubated at 37 °C under constant light for 2 days. Colony-forming units (CFUs) were counted manually. All assays were performed in three independent biological replicates, with three technical replicates per sample.

Spore viability was calculated as the percentage of viable conidia relative to the initial fresh spore inoculum, using the formula: Viability (%) = (Number of CFUs after treatment / Number of CFUs from untreated fresh spores) × 100

Statistical significance was determined using unpaired two-tailed Student’s t-tests, comparing the mean values of indicated mutant strains to the wild-type control. Data are presented as mean ± standard deviation (SD) of three independent experiments.

### Microscopy

Representative colonies of *A. nidulans* were imaged using an SZX12-ILLB2-200 stereomicroscope (Olympus Corporation, Tokyo, Japan). Digital images were captured with an Olympus SC30 digital camera coupled to the stereomicroscope, and post-acquisition image processing was performed using cellSens software (Olympus).

### Structural modeling and analysis of VelB and related variants

Three-dimensional structures of VelB and its derived variants VelB^R71A^, VelB^R80A^, and VelB^R81A^ were predicted using AlphaFold 3 (Abramson et al. 2024). The crystal structure of the *A. nidulans* VosA-VelB heterodimer (PDB ID: 4N6R) (Ahmed et al. 2013) was employed as the reference structural template for modeling. The highest-confidence model generated by AlphaFold 3 was used for further analysis. The electrostatic potential at each atomic position was calculated using the Coulomb Potential model (Honig and Nicholls 1995).

## Results

### Sequence features of velvet DNA-binding region and alanine scanning of conserved residues by *A. nidulans* VelB

*In silico* analysis of amino acid conservation within the velvet DNA-binding domain was performed using 4,999 putative velvet domains derived from diverse fungal species, and the corresponding sequence logo was constructed (**Figure 1A**). The results revealed that the velvet DNA-binding domain consists of approximately 30 amino acid residues, among which multiple residues exhibit strong evolutionary conservation. Notably, several positively charged amino acids were presumed to be functionally critical.

**Figure 1.**
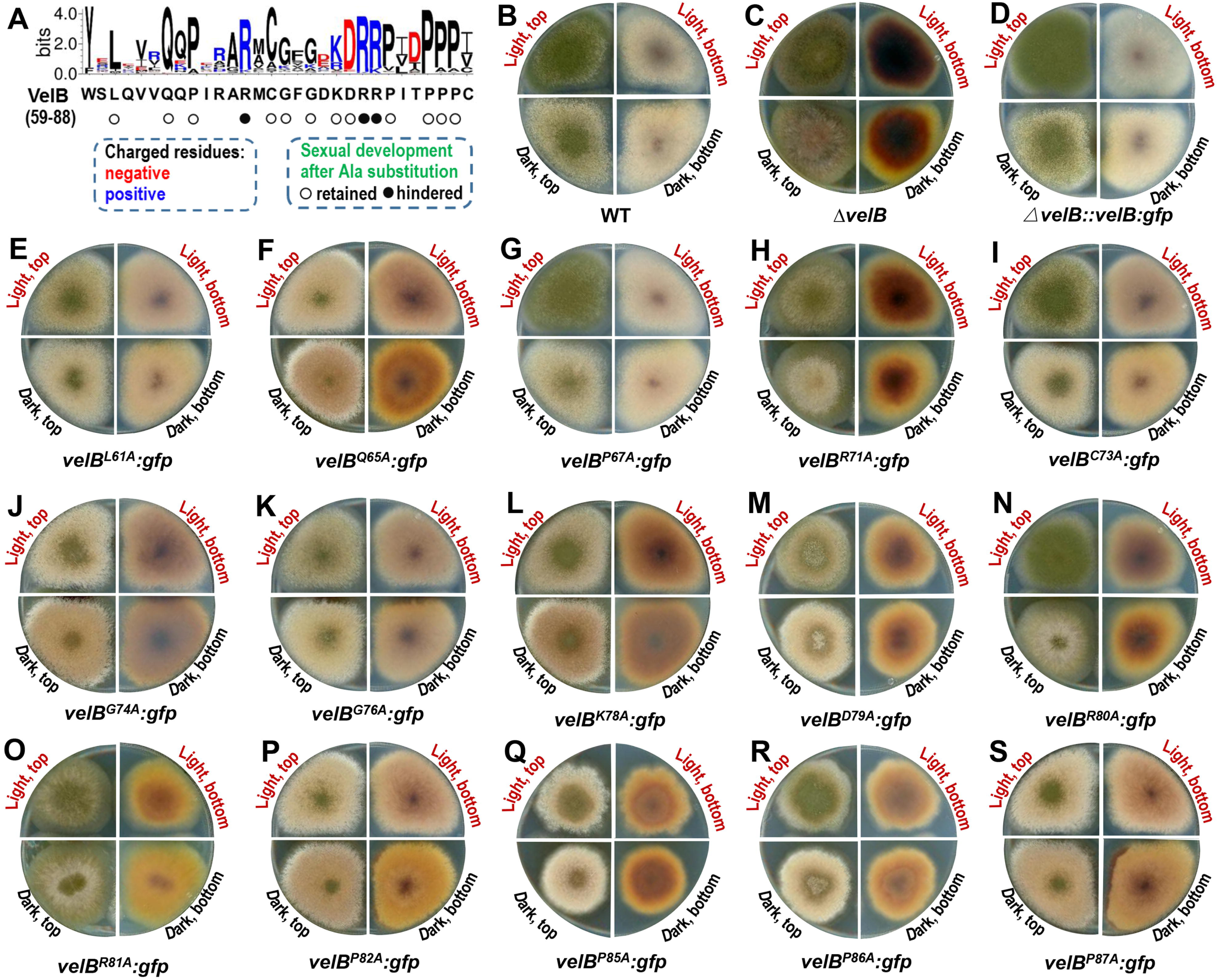
Alanine scanning of 15 conserved residues in the velvet DNA-binding region of *A. nidulans* VelB. A. Sequence logo of 4,999 velvet DNA-binding regions and mutagenesis summary. Residues are colored by charge (blue: positive; red: negative; black: neutral). B–S. Colony morphology of strains cultured on MM plates at 37°C for 5 days. B-D respectively correspond to the wild type (WT, AGB551), *velB* knockout (AGB1064), and *velB* complement (AGB1192), which were used as comparison with the mutants. E-S respectively correspond to *velB^L61A^:gfp* (AGB1484), *velB^Q65A^:gfp* (AGB1485), *velB^P67A^:gfp* (AGB1486), *velB^R71A^:gfp* (AGB1487), *velB^C73A^:gfp* (AGB1488), *velB^G74A^:gfp* (AGB1489), *velB^G76A^:gfp* (AGB1490), *velB^K78A^:gfp* (AGB1491), *velB^D79A^:gfp* (AGB1492), *velB^R80A^:gfp* (AGB1493), *velB^R81A^:gfp* (AGB1494), *velB^P82A^:gfp* (AGB1495), *velB^P85A^:gfp* (AGB1496), *velB^P86A^:gfp* (AGB1497), and *velB^P87A^:gfp* (AGB1498).

To identify the key residues governing the biological functions of velvet proteins, alanine-scanning mutagenesis on 15 conserved residues (conservation score > 1.5 bits) in *A. nidulans* VelB was conducted, wherein each target residue was individually substituted with alanine. *A. nidulans* VelB was selected as the ideal model for investigating the structure-function relationship of velvet family proteins for two reasons: first, of the four velvet homologs in A. nidulans, VelB features the shortest amino acid sequence and the highest cross-species conservation across fungi (Chen et al. 2024); second, functional perturbation of VelB induces distinct phenotypic alterations (Chen et al. 2025). The colonial phenotypes of all mutant strains were compared with those of the wild-type (WT), *velB* knockout and complementary strains (**Figure 1B–S**). Preliminary phenotypic characterization indicated that three mutants, namely *velB^R71A^*, *velB^R80A^*and *velB^R81A^*, displayed the most dramatic phenotypic defects.

### Arginine 71, 80 and 81 of the VelB DNA-binding region are essential for spore viability, sexual development and secondary metabolism

The impacts of the *velB^R71A^*, *velB^R80A^*, and *velB^R81A^* mutations on fungal development and secondary metabolites were further investigated (**Figure 2**). Under asexual growth condition (in light), *A. nidulans* WT and *velB* complement produced green conidia, whereas the *velB*-mutated strains yielded grey-green conidia (**Figure 2A**). Under sexual growth conditions (in dark and sealed), *A. nidulans* WT and *velB* complement favors cleistothecia formation, however, the mutated strains *velB^R80A^* and *velB^R81A^*failed to generate cleistothecia on MM, a phenotype identical to that of the *velB* knockout strain (**Figure 2A**). The *velB^R71A^* mutant exhibited severely impaired cleistothecia production, with the yield accounting for only 13.3% of that in the WT strain.

**Figure 2.**
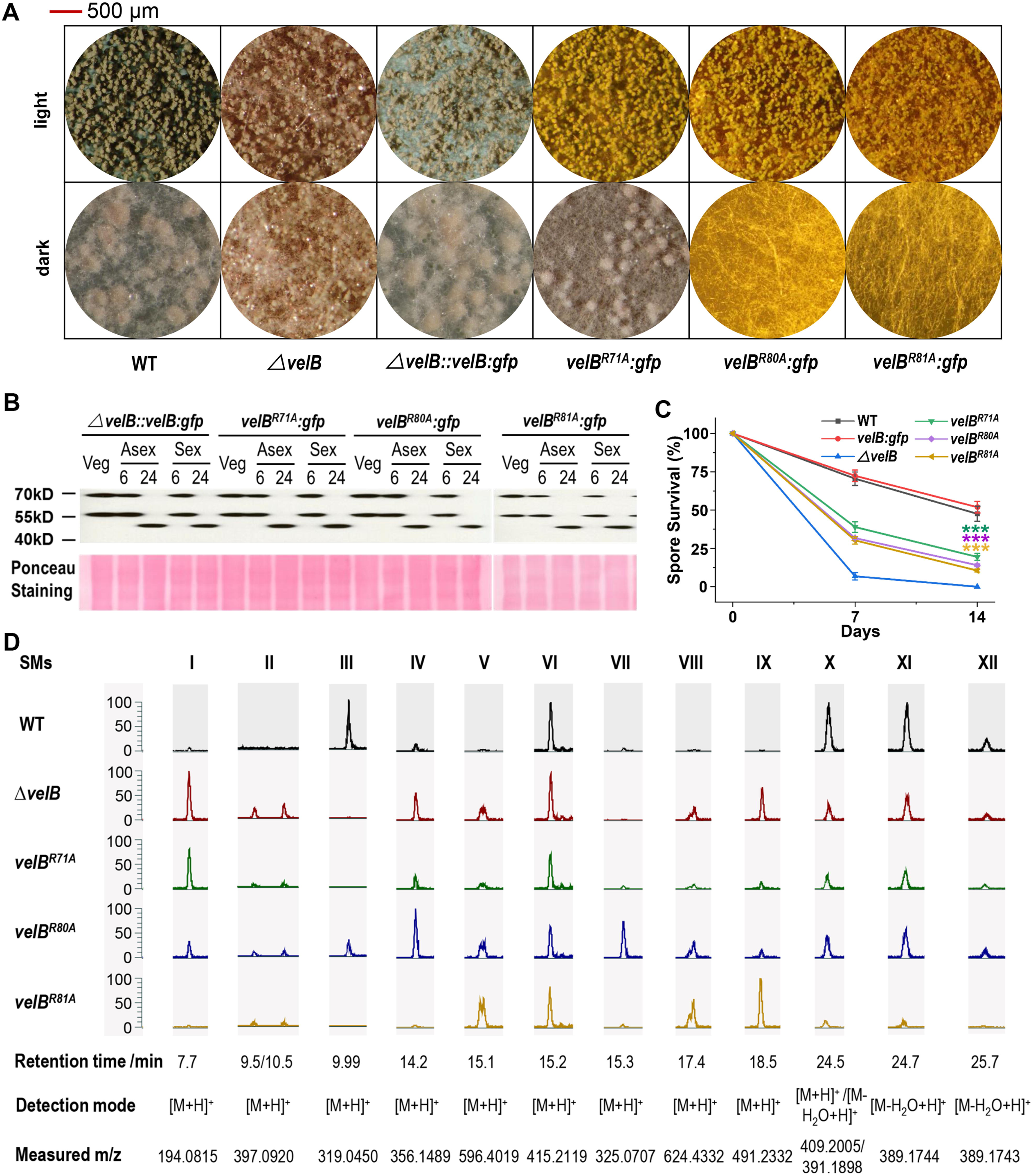
Point mutations of R71A, R80A or R81A disrupt VelB function and cause defective fungal development in *A. nidulans*. A. Fungal developmental phenotypes under asexual (light) and sexual (dark) conditions. The wild type (WT), the *velB* deletion strain, and the VelB-green fluorescent protein (GFP) complementation strain served as controls. Colony morphology was documented on day 5 post-inoculation. B. Western blot analysis of VelB^R71A^-GFP, VelB^R80A^-GFP and VelB^R81A^-GFP expression patterns under sexual, asexual, and vegetative growth conditions. The VelB-GFP strain was used as a control. For all experimental conditions, spores were incubated overnight in liquid MM. For sexual and asexual inductions, mycelia were transferred to solid MM for 6 or 24 h under sexual and asexual conditions, respectively. Ponceau staining was used as a loading control for protein normalization. The VelB-GFP Western blot panel was adapted from Figure 2C of the previous study (Chen et al. 2025) under the Creative Commons Attribution 4.0 International license. C. Conidial viability assays in the WT and *velB* mutant strains and complement. 100 conidia per strain were plated after zero, seven and 14 d treatment, with initial colony-forming units set to 100%. Error bars represent standard error of the mean from n ≥ 3 independent biological replicates (***, *P*<0.001). D. SM profiles of WT and *velB* mutant strains. Extracted ion chromatograms of 12 detected SMs are shown. The numbers indicate the identified SMs: I, cichorine (Sanchez et al. 2012); II, asperthecin (Szewczyk et al. 2008); III, F9775A/B (Sanchez et al. 2010); IV, arguosin H (Nielsen et al. 2011); V, emericellamide C (Chiang et al. 2008); VI, austinoneol A (Lo et al. 2012); VII, sterigmatocystin (Yu and Leonard 1995); VIII, emericellamide E (Chiang et al. 2008); IX, terrequinone A (Balibar et al. 2007; Bouhired et al. 2007); X, emericellin (Sanchez et al. 2011); XI, shamixanthone (Sanchez et al. 2011); and XII, epishamixanthone (Sanchez et al. 2011). The chromatogram panels of WT and *velB* knockout strains were adapted from Figure 3B of the previous study (Chen et al. 2025) under the Creative Commons Attribution 4.0 International license.

The protein expression profiles were compared by Western blotting experiments and the results revealed that compared to the *velB:gfp* expressing strain, the mutated VelBs were properly expressed (**Figure 2B**). The full-length VelB-GFP protein with a molecular weight of 67 kD was consistently detected across vegetative growth stages, as well as the early developmental stages of both asexual and sexual reproduction. Furthermore, long-term conidial viability was assessed among the WT and *velB* mutants and complement strain on solid MM (**Figure 2C**). Conidia of the mutated strains *velB^R71A^*, *velB^R80A^*, and *velB^R81A^* exhibited a rapid loss in viability relative to WT conidia, after seven days and thereafter. In particular, on the 14th day, the WT and complemented strains maintained a conidial survival rate of approximately 50%. In contrast, the three point-mutant strains displayed a statistically significant reduction in spore viability, with survival rates dropping to roughly 15%. However, unlike the *velB* knockout strain, which completely lost conidial viability after 14 days treatment, the *velB^R71A^*, *velB^R80A^*, and *velB^R81A^* mutants retained partial conidial viability under identical conditions.

The accumulation patterns of 12 secondary metabolites (SMs) were compared among the *A. nidulans* WT and *velB* mutant strains (**Figure 2D**). The *velB* variants exhibited disrupted secondary metabolism, and their metabolic profiles partially overlapped with those of the *velB* deletion strain.

Compared with the WT, both *velB* variants and deletion strain showed attenuated or abolished biosynthesis of emericellin, epishamixanthone, F9775A/B, as well as shamixanthone. By contrast, the production of asperthecin, emericellamide C, emericellamide E, and terrequinone A was significantly upregulated in these mutant strains. Additionally, austinoneol A displayed stable accumulation levels across all tested WT and *velB* mutant strains. Collectively, these multiple lines of evidence demonstrate that R71, R80, and R81 serve as pivotal functional residues of VelB, and are essential for modulating fungal development and secondary metabolism in *A. nidulans*.

### Point mutations of R71A, R80A and R81A trigger local reduction and global remodeling of electrostatic potential

Three positively charged arginine residues (R71, R80 and R81) of VelB DNA-binding region contribute substantial positive electrostatic potential, which is essential for mediating electrostatic attraction between the protein and the negatively charged DNA molecule. To systematically characterize how mutations at these sites alter the surface electrostatic potential, electrostatic calculations were performed based on the resolved crystal structures of wild-type VelB and its mutant variants. A comparative analysis was conducted (**Figure 3**), and core statistical data regarding electrostatic features are summarized in **Table 2**.

**Figure 3.**
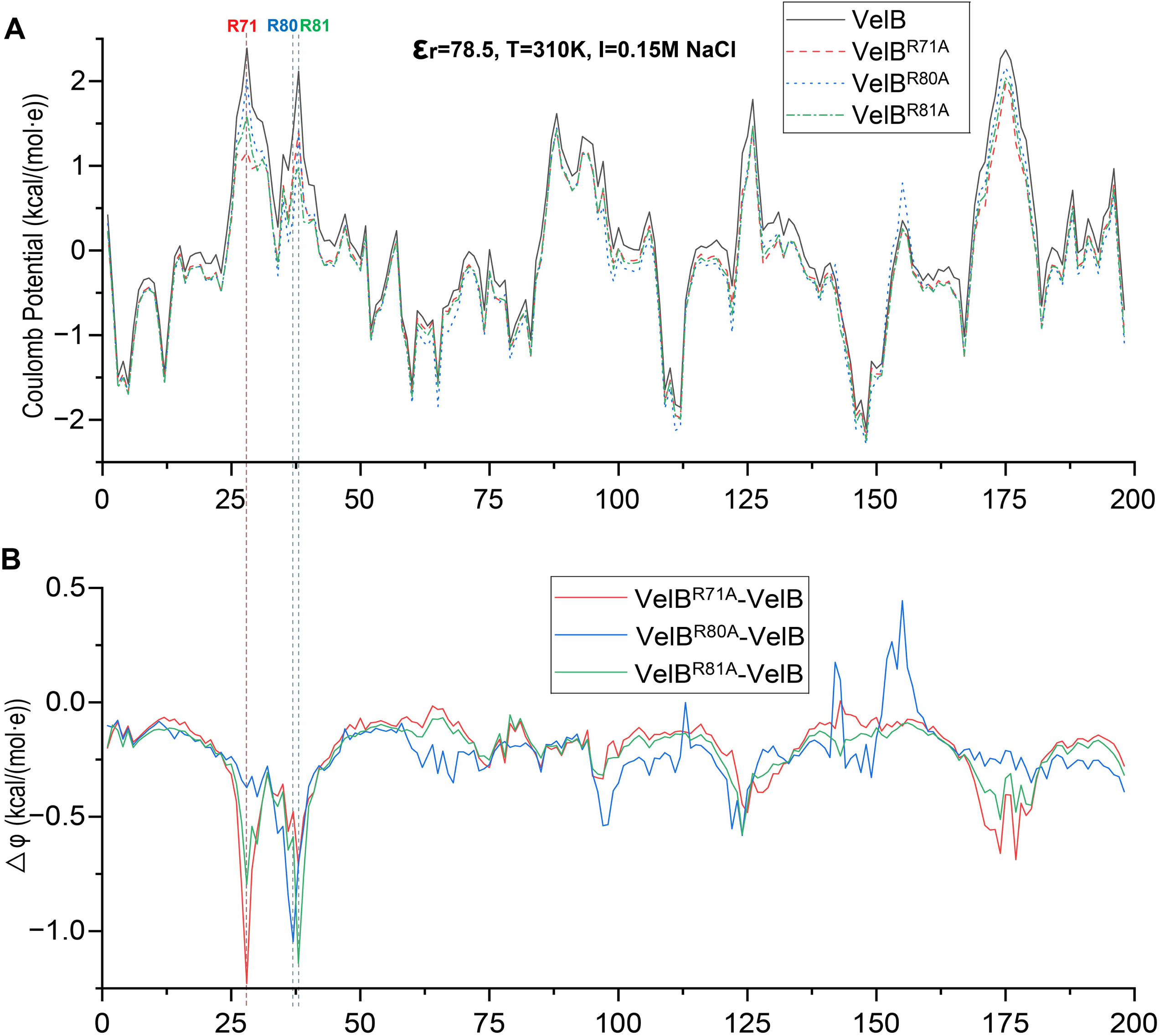
Electrostatic potential dynamics pre- and post-mutation. A. Coulomb potentials of VelB, VelB^R71A^, VelB^R80A^, and VelB^R81A^ at each residue. The truncated VelB with 198 residues from the *A. nidulans* VosA-VelB heterodimer (PDB ID: 4N6R) (Ahmed et al. 2013) was employed as the reference structural template for modeling. The R71, R80, and R81 correspond to R28, R37, and R38 of the truncated VelB as indicated in the figure. B. Electrostatic potential changes of VelB^R71A^, VelB^R80A^, and VelB^R81A^ relative to VelB. The value shown at each residue site in the figure represents the difference in electrostatic potential between the mutant VelB and the original VelB. A larger absolute value indicates a greater magnitude of change.

**Table 2.** Key Electrostatic Statistics among VelB and its variants.

| Parameters | VelB | VelB<sup>R71A</sup> | VelB<sup>R80A</sup> | VelB<sup>R81A</sup> |
| --- | --- | --- | --- | --- |
| Coulomb mean<br>(kcal/(mol*e)) | +0.0477 | -0.1673 | -0.1818 | -0.1795 |
| Coulomb<br>minimum | -2.106 | -2.216 | -2.290 | -2.246 |
| Coulomb<br>maximum | +2.391 | +1.965 | +2.158 | +2.037 |
| Mutation Site/<br>Original<br>Potentials |  | +1.162/+2.391 | +0.391/+1.437 | +0.976/+2.116 |
| Residues with<br>$\vert \Delta\phi\vert > 0.01$ | | 197/198<br>(99.5%) | 197/198<br>(99.5%) | 198/198<br>(100%) |
| Residues with<br>$\vert \Delta\phi\vert > 0.05$ | | 194/198<br>(98.0%) | 196/198<br>(98.9%) | 198/198<br>(100%) |
| Residues with<br>$\vert \Delta\phi\vert > 0.10$ | | 155/198<br>(78.3%) | 189/198<br>(95.5%) | 188/198<br>(94.9%) |
| Maximum $\vert \Delta\phi\vert $ | | 1.229<br>(Mutation<br>Site) | 1.046<br>(Mutation<br>Site) | 1.140 (Mutation<br>Site) |

The analysis revealed that compared to the original VelB, all three mutations significantly decreased the mean, minimum and maximum Coulomb potential values of the entire protein. R71 is positioned at the strongest electropositive region of VelB, with an original potential of +2.391 kcal/(mol·e). Alanine substitution at this site (R71A) resulted in the most dramatic reduction in electrostatic potential, with an absolute decrease of 1.229 kcal/(mol·e), demonstrating that R71 is a critical residue for maintaining the strong positive electrostatic environment. Differently, R81 showed the weakest intrinsic positive potential (+1.44 kcal/(mol·e)) among the three arginines. We speculate that this is caused by strong local electrostatic shielding from the nearby negatively charged aspartate residues D77 and D79. Beyond local changes, each mutation triggers extensive global remodeling of the protein electrostatic profile. Further residue-level analysis confirmed that the resultant electrostatic disturbances propagate widely, affecting nearly every amino acid site across the protein structure.

### Replacement of the VelB DNA-binding region by a *Capsaspora* velvet one

The velvet protein from *Capsaspora owczarzaki* represents the first identified velvet-family member outside the fungal kingdom (Ahmed et al. 2013). To explore the functional compatibility of the *Capsaspora* velvet DNA-binding region in *A.nidulans* VelB, heterologous domain replacement assays was conducted (**Figure 4**). Sequence alignment between *Capsaspora* velvet DNA-binding region and four *A. nidulans* ones revealed that the former lacks the positively charged arginine residue corresponding to VelB Arg71 (**Figure 4A**). Phenotypic assays demonstrated that the chimeric strain *velB*(Δ*velB_DBR::CapVelvet_DBR*), in which the native VelB DNA-binding region was substituted with its *Capsaspora* counterpart, failed to fully rescue the phenotypic defects of the *velB* deletion strain. This chimeric mutant only exhibited partial recovery of sexual development under dark culture conditions (**Figure 4B**), showing a phenotypic profile highly analogous to that of the mutant *velB^R71A^* described above (**Figure 2A**). To verify the indispensable role of R71, this key residue was artificially introduced into the *Capsaspora* velvet DNA-binding region to generate the modified chimeric strain *velB (*Δ*velB_DBR::CapVelvet_DBR)^R71^*. Subsequent phenotypic analyses revealed that supplementation with R71 substantially restored the defective phenotypes of the *velB* knockout strain (**Figure 4B**). Notably, all mutants involved in this assay were constructed based on *velB* cDNA to eliminate potential splicing interference. Western blotting was subsequently performed to verify protein expression, and the results confirmed that all chimeric VelB variants were successfully expressed at normal patterns relative to the control *velBcDNA:gfp* expressing strain (**Figure 4C**).

**Figure 4.**
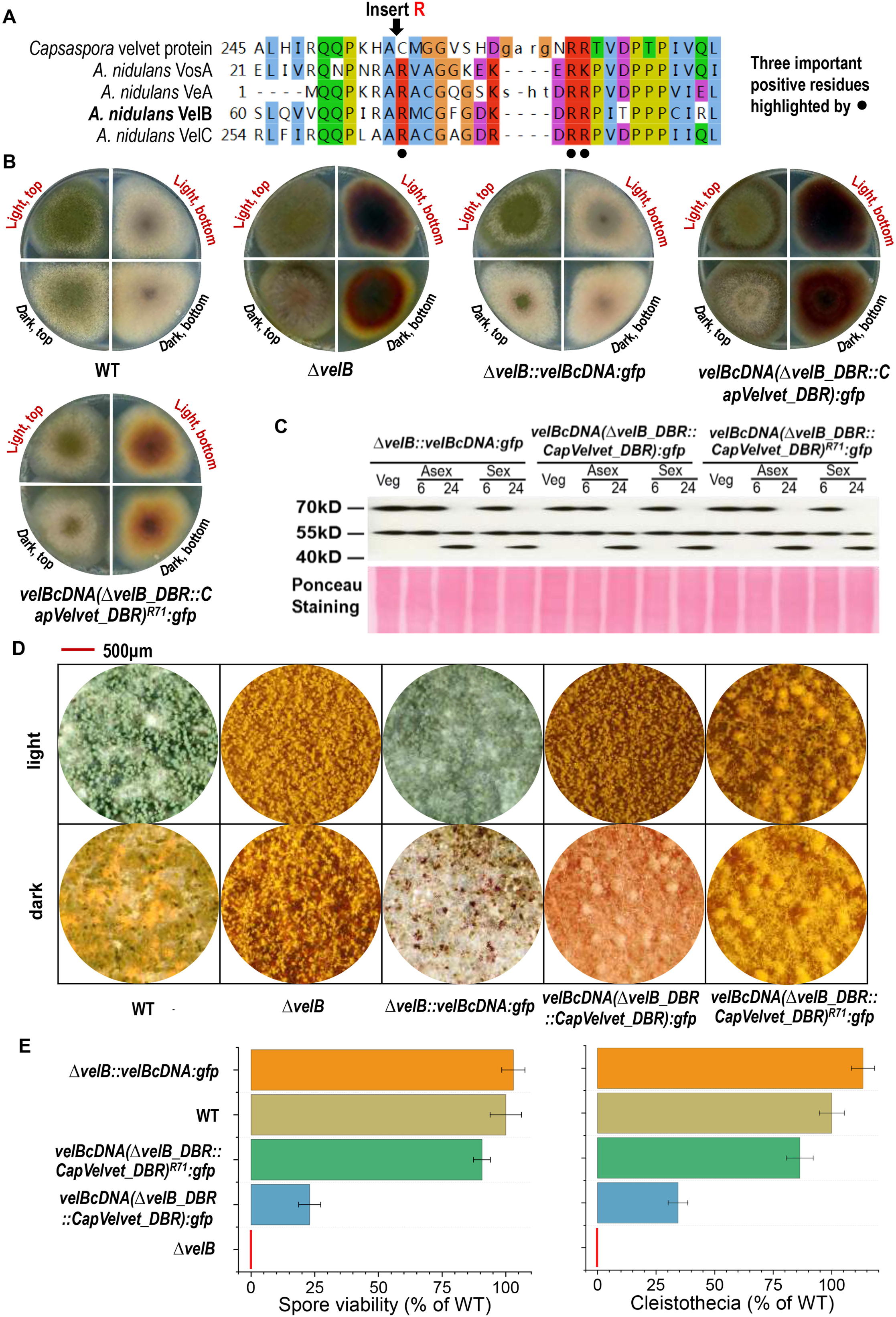
Replacement of *A. nidulans* VelB DNA-binding region (DBR) by *Capsaspora* velvet (CapVelvet) one. A. Multiple sequence alignment of DNA-binding regions across *Capsaspora* velvet protein, *A. nidulans* VosA, VeA, VelB and VelC. Solid black balls denote the three crucial positively charged residues as revealed in Figure 2A. B. Colony phenotypic comparison of *A. nidulans* WT, *velB* deletion mutant (Δ*velB*), VelB-GFP complemented strain (expressing C-terminally GFP-tagged *velB* cDNA in the Δ*velB* background), *velBcDNA(*Δ*velB_DBR::CapVelvet_DBR):gfp*, and *velBcDNA(*Δ*velB_DBR::CapVelvet_DBR)^R71^:gfp*. All strains were grown on solid MM under asexual (light) or sexual (dark, sealed) inducing conditions for five days post-inoculation. All mutants were generated using the *A. nidulans velB* cDNA backbone. Notably, the native Capsaspora velvet DBR lacks one of the three conserved positively charged residues present in VelB; an arginine residue (R71) was therefore inserted to restore this position, as indicated in Figure 4A. C. Western blot analysis of VelB-GFP and its chimeric/mutant derivatives, all expressed from the native *velB* promoter. Protein detection was performed using an anti-GFP antibody, with Ponceau staining serving as a loading control for protein normalization. Protein abundance was assessed at multiple developmental stages: vegetative growth (24 h submerged culture), asexual development (6 h and 24 h light-exposed plates), and sexual development (6 h and 24 h dark-incubated plates), all conducted at 37°C. The predicted molecular mass of all full-length fusion proteins is approximately 67 kDa. D. Representative micrographs of colonies on day 5 post-inoculation. A red scale bar is provided for size reference. E. Conidial viability assays and cleistothecial quantification. Conidial viability of the indicated strains was tested after 14 days treatment, with viability of the WT strain set to 100%. Cleistothecia of indicated strains were quantified from plated cultures grown under sexual inducing conditions for 5 d post- inoculation, with cleistothecial production by the WT strain set to 100%. Error bars represent standard error of the mean from n ≥ 3 independent biological replicates.

The phenotypic characteristics of the two chimeric strains, *velBcDNA(*Δ*velB_DBR::CapVelvet_DBR)*, and *velBcDNA(*Δ*velB_DBR::CapVelvet_DBR)^R71^*, in fungal development and secondary metabolism were further investigated. Under dark sealed culture conditions for sexual induction, both chimeric mutants were capable of forming cleistothecia on MM (**Figure 4D**). The chimeric strain *velBcDNA(*Δ*velB_DBR::CapVelvet_DBR)^R71^*also significantly reduced colony pigmentation levels. The long-term spore viability after 14 days treatment and cleistothecia amounts were compared among the WT and *velB* mutants and complemented strain on solid MM (**Figure 4E**). For standardized quantitative comparison, spore viability rate and Cleistothecia amount of the WT strain were normalized to 100%, whereas those of *velB* knockout were set to 0% due to its absence of long-term spore viability and Cleistothecia formation. Quantitative data showed that the *velBcDNA(*Δ*velB_DBR::CapVelvet_DBR)* mutant exhibited partial phenotypic restoration, with a 14-day conidial survival rate of approximately 23% and a cleistothecial abundance of 34% relative to the WT. Importantly, the additional introduction of the positively charged residue R71 dramatically enhanced the biological functionality of the chimeric protein. The modified *velB*(Δ*velB_DBR::CapVelvet_DBR*) mutant restored conidial viability to 91% and cleistothecial formation to 86% of the WT level, highlighting the essential function of R71 in empowering the conserved DNA-binding region for VelB-mediated physiological regulation.

### Positive residues are frequently present in various DNA-binding regions

There are various transcription factor proteins harboring different types of DNA-binding regions. To examine whether positively charged residues are broadly conserved and prevalent across different DNA-binding regions, their sequence features were investigated using the InterPro database (https://www.ebi.ac.uk/interpro). The analysis revealed that positively charged residues are widely distributed across diverse DNA-binding regions and typically exhibit high evolutionary conservation. **Figure 5** shows the sequence logos of six representative DNA-binding regions from Rel, Zinc finger, homobox, basic region leucine zipper, serum response factor, and helix-loop-helix transcription factors.

**Figure 5.**
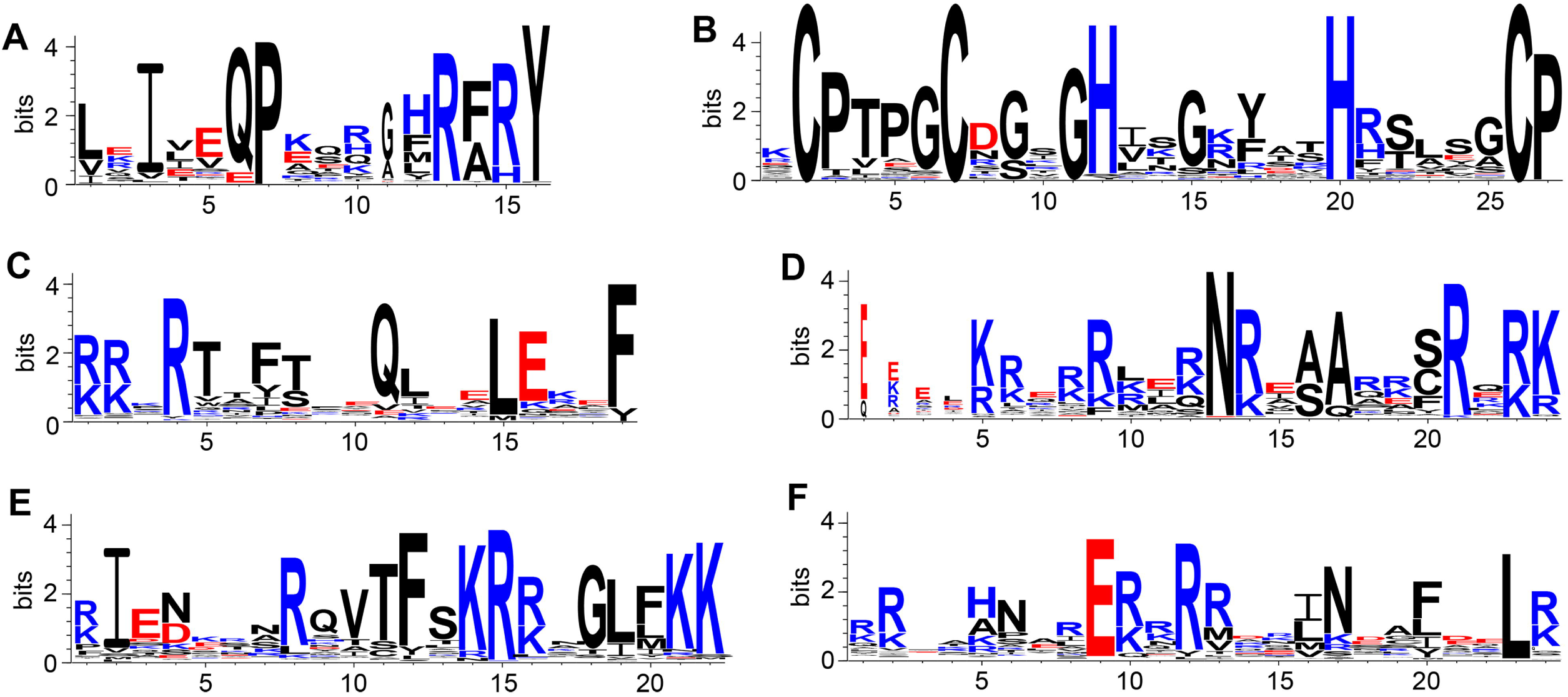
Sequence logo of representative DNA-binding regions. Residues are colored by charge that positive ones are in blue, negative ones are in red, and neutral ones are in black. A to F correspond to DNA-binding domains of Rel (PF00554), Zinc finger (PF01530), homobox (PF00046), basic region leucine zipper (PF00170), serum response factor (PF00319), and helix-loop-helix (PF00010) transcription factors, respectively.

## Discussion

The velvet family of transcriptional and epigenetic regulatory proteins represents an ancient class of DNA-binding proteins, with orthologs traceable to the pre-fungal ancestor that predated the divergence of the opisthokont lineage (Chen et al. 2024). Velvet regulators have attracted broad attention from fungal scientists due to their wide distribution in the fungal kingdom, and unique and crucial roles in fungal development, secondary metabolism, stress response, and pathogenicity (Chen et al. 2026). Recently, the general architecture of velvet domains was proposed by dividing them into an approximately 30-amino-acid N-terminal DNA-binding region and a roughly 100-amino-acid C-terminal dimerization region, which markedly clarifies the working mechanisms of velvet proteins (Chen et al. 2025).

In the present work, the DNA-binding region of velvet proteins has been systemically characterized by using *A. nidulans* VelB as a paradigm. Through alanine scanning mutagenesis targeting 15 conserved amino acid residues within this DNA-binding region of *A. nidulans* VelB, three key positively charged residues, namely R71, R80 and R81, were identified (**Figure 1**). Subsequent functional assays demonstrated that these three residues are essential for the full function of VelB, including the maintenance of long-term spore viability, esiduesregulation of sexual development and modulation of secondary metabolism (**Figure 2**). Structural analysis of the VosA homodimer and VosA-VelB heterodimer revealed a strongly electropositive surface across the velvet DNA-binding region and these three corresponding positively charged residues in VosA were verified their important roles in DNA-binding (Ahmed et al. 2013). To clarify the effects of the above mutations on surface electrostatic potential, the electrostatic profiles of wild-type VelB and its mutant variants were compared. The results showed that individual single-point mutations not only significantly lower local electropositive potential but also provoke long-range electrostatic perturbations, ultimately altering the global surface profile critical to molecular recognition and protein functionality (**Figure 3** and **Table 2**).

These three positions, which are invariably occupied by positively charged (basic) residues, exhibit strict evolutionary conservation across velvet domains from all major taxonomic groups. Frequency analysis of 4,999 non-redundant velvet domain sequences revealed that position 71 is occupied by R in 85% of sequences and K in 4%, position 80 by R in 91% and K in 2%, and position 81 by R in 84% and K in 9% (**Figure 1A**). To test the functional conservation of the velvet DNA-binding domain, a cross-kingdom domain-swapping experiment was performed, in which the endogenous DNA-binding region of *A. nidulans* VelB was replaced with the orthologous region from *C. owczarzaki*—a unicellular opisthokont that diverged prior to the emergence of the fungal kingdom (**Figure 4**). However, expression of this chimeric VelB construct in a *velB* deletion strain failed to rescue the full spectrum of *velB* mutant phenotypes, with only partial restoration of sexual development observed under dark conditions. Sequence alignment revealed that the *Capsaspora* velvet domain specifically lacks the critical arginine residue at the position corresponding to R71 in VelB (**Figure 4A**), which is essential for VelB function (**Figure 2**). To directly test whether this single residue difference accounts for the functional incompatibility, an arginine residue at the corresponding position in the *Capsaspora* velvet DNA-binding region was artificially introduced to generate the modified chimeric strain *velB (*Δ*velB_DBR::CapVelvet_DBR)^R71^*. Strikingly, expression of this modified chimeric protein in the *velB* deletion strain was sufficient to substantially rescue all tested mutant phenotypes, including long-term spore viability, sexual development and colony pigmentation (**Figure 4B, D, & E**). These findings strongly suggest that while the high sequence conservation of the velvet DNA-binding domain underpins a broadly conserved core mechanism of protein-DNA interaction, subtle functional divergence may have emerged among distinct velvet family members.

In summary, using *A. nidulans* VelB as a paradigm, we have identified three critical positively charged residues within the velvet DNA-binding region that are indispensable for VelB function. These residues constitute the structural prerequisites for generating the strongly electropositive surface potential that mediates specific interactions between velvet proteins and DNA. Our findings not only advance the mechanistic understanding of fungal velvet regulators, but also provide general insights into the molecular basis of DNA recognition. Indeed, surface-exposed positively charged residues represent a universal feature of diverse DNA-binding motifs, including those of Rel homology domain, basic region leucine zipper and helix-loop-helix transcription factors (**Figure 5**).

## Acknowledgments

WC was supported by Humboldt fellowship during his stay in Göttingen.

