## Supplementary material for "Characterization of velvet DNA-binding region by *Aspergillus nidulans* VelB": Table S1; Table S2

### Materials and Methods

#### Plasmid construction and preparation

For plasmid construction, target DNA fragments were amplified by polymerase chain reaction (PCR). The amplified fragments were ligated into the designated restriction sites of plasmids using the Seamless Cloning and Assembly Kit (Invitrogen). Plasmids were then propagated in *E. coli* and extracted using the Qiaprep Spin Miniprep Kit (Qiagen) in accordance with the manufacturer's instructions. All plasmids generated and employed in this work are listed in Table S1.

**Table S1. Plasmids constructed and used in this study.** natRM = nourseothricin recyclable marker cassette, phleoRM = phleomycin recyclable marker cassette

| Plasmid | Description | Source |
|---|---|---|
| pME4319 | The recyclable marker cassette-containing plasmid with the bleo gene conferring resistance to phleomycin | (Liu et al. 2021) |
| pME4687 | *A. nidulans velB* with sgfp tag: velB::sgfp::natRM | (Kohler et al. 2026) |
| pME5332 | *A. nidulans velB* 3UTR integrated in the PmlI restriction cutting site of plasmid pME4319 | (Chen et al. 2025) |
| pME5333 | *A. nidulans* velB_5UTR:velBcDNA: sgfp: phleoRM:velB_3UTR | (Chen et al. 2025) |
| pME5335 | *A. nidulans* velB_5UTR: velBcDNA (ΔvelB_DBR::CapVelvet_DBR): sgfp: phleoRM: velB_3UTR | This study |
| pME5338 | *A. nidulans* velB_5UTR: velB$^{L61A}$:sgfp: phleoRM: velB_3UTR | This study |
| pME5339 | *A. nidulans* velB_5UTR: velB$^{Q65A}$:sgfp: phleoRM: velB_3UTR | This study |
| pME5340 | *A. nidulans* velB_5UTR: velB$^{P67A}$:sgfp: phleoRM: velB_3UTR | This study |
| pME5341 | *A. nidulans* velB_5UTR: velB$^{R71A}$:sgfp: phleoRM: velB_3UTR | This study |
| pME5342 | *A. nidulans* velB_5UTR: velB$^{C73A}$:sgfp: phleoRM: velB_3UTR | This study |
| pME5343 | *A. nidulans* velB_5UTR: velB$^{G74A}$:sgfp: phleoRM: velB_3UTR | This study |
| pME5344 | *A. nidulans* velB_5UTR: velB$^{G76A}$:sgfp: phleoRM: velB_3UTR | This study |
| pME5345 | *A. nidulans* velB_5UTR: velB$^{K78A}$:sgfp: phleoRM: velB_3UTR | This study |
| pME5346 | *A. nidulans* velB_5UTR: velB$^{D79A}$:sgfp: | This study |

| | | |
|---|---|---|
| | phleoRM: velB_3UTR | |
| pME5347 | *A. nidulans* velB_5UTR: velB$^{R80A}$:sgfp: phleoRM: velB_3UTR | This study |
| pME5348 | *A. nidulans* velB_5UTR: velB$^{R81A}$:sgfp: phleoRM: velB_3UTR | This study |
| pME5349 | *A. nidulans* velB_5UTR: velB$^{P82A}$:sgfp: phleoRM: velB_3UTR | This study |
| pME5350 | *A. nidulans* velB_5UTR: velB$^{P85A}$:sgfp: phleoRM: velB_3UTR | This study |
| pME5351 | *A. nidulans* velB_5UTR: velB$^{P86A}$:sgfp: phleoRM: velB_3UTR | This study |
| pME5352 | *A. nidulans* velB_5UTR: velB$^{P87A}$:sgfp: phleoRM: velB_3UTR | This study |
| pME5353 | *A. nidulans* velB_5UTR: velBcDNA (ΔvelB_DBR:: CapVelvet_DBR)$^{R71}$:sgfp: phleoRM: velB_3UTR | This study |

**Oligonucleotides used in this study**

Oligonucleotides used in this study are listed in Table S2.

**Table S2. Oligonucleotides utilized in this study**

| Oligo Name | Sequence (5'-3') |
|---|---|
| WC12 | ATAGGCCTGAGATTTCTACTTGTACAGTTCGTCCAT |
| WC30 | TGCAGGAATTCGATATTTGTTTAAAC |
| WC154 | ACACCACCCATGCAGGCGTGCTTGGGCTGCTGGCGGATGTG CAAGGACCAGATACGACC |
| WC155 | CCTGCATGGGTGGTGTCTCCCACGACGGTGCCCGCGGTAAC CGAAGGCCAATCACACCC |
| WC171 | CACGACTTGCAAGGACCAGAT |
| WC172 | GTCCTTGCAAGTCGTGGCACAGCCGATTCGCGC |
| WC173 | CTGTTGCACGACTTGCAAGG |
| WC174 | GCAAGTCGTGCAACAGGCGATTCGCGCGCGGA |
| WC175 | CGCGCGAATCGGCTGTT |
| WC176 | ACAGCCGATTCGCGCGGCGATGTGCGGCTTTGG |
| WC177 | CGCACATCCGCGCGCGAAT |
| WC178 | CGCGCGCGGATGTGCGCCTTTGGCGATAAGGT |
| WC179 | ATCGCCAAAGCCGCACATC |
| WC180 | GTGCGGCTTTGGCGATGCGGTTGGCTATTTGTA |
| WC181 | CCTAGAGAGAGTCAGCACGGAC |
| WC182 | GCTGACTCTCTCTAGGCCCGAAGGCCAATCAC |
| WC183 | GTCCTAGAGAGAGTCAGCACGG |
| WC184 | TGACTCTCTCTAGGACGCAAGGCCAATCACACC |

| | |
|---|---|
| WC185 | TCGGTCCTAGAGAGAGTCAGCA |
| WC186 | CTCTCTCTAGGACCGAGCGCCAATCACACCCCC |
| WC187 | CCTTCGGTCCTAGAGAGAGTCA |
| WC188 | TCTCTAGGACCGAAGGGCAATCACACCCCCTC |
| WC189 | TGTGATTGGCCTTCGGTCCT |
| WC190 | CCGAAGGCCAATCACAGCCCCTCCGTGTATCC |
| WC191 | GGGTGTGATTGGCCTTCGGT |
| WC192 | AAGGCCAATCACACCCGCTCCGTGTATCCGTC |
| WC193 | AGGGGGTGTGATTGGCCTT |
| WC194 | GCCAATCACACCCCCTGCGTGTATCCGTCTAA |
| WC208 | GGACCAGATACGACCGTCATG |
| WC209 | CGGTCGTATCTGGTCCGCGCAAGTCGTGCAACA |
| WC210 | CAAAGCCGCACATCCGCGCG |
| WC211 | CGGATGTGCGGCTTTGCCGATAAGGTTGGCTA |
| WC212 | CATGCAACGGGCGTGCTTGGGCTGCTGGC |
| WC213 | GCACGCCCGTTGCATGGGTGGTGTCTCCCA |
| WC251 | CATCCGCGCGCGAATCGGCTGTT |
| WC252 | GATTCGCGCGCGGATGGCCGGCTTTGGCGATAA |

Note: the underlined oligonucleotides were generated the overlaps for seamless cloning in plasmid construction.

https:// 10.7554/eLife.68058.
